# Phytoplankton recruitment of specific microbial assemblages and phylosymbiotic patterns

**DOI:** 10.1101/2025.02.10.637405

**Authors:** Joo-Hwan Kim, Jin Ho Kim, Zhun Li, Hyeon Ho Shin, Young Kyun Lim, Seung Ho Baek, Pengbin Wang, Ro Young Park, Soo Hwan Jin, Myung-Soo Han, Bum Soo Park

## Abstract

Phytoplankton and bacteria represent two major pillars of the carbon cycle in marine ecosystems. While their interactions are known to be tightly linked, the specific mechanisms underlying these interactions remain largely unexplored. Evidence of host-specific microbial assemblages could serve as a foundation for studies of detailed host-microbe interactions, yet such research remains limited in phytoplankton. Here, we not only investigate the microbial assemblages of six phytoplankton species, including multiple strains of dinoflagellates and diatoms, but also samples of phytoplankton blooms from the field. Our results reveal the presence of host-specific microbial assemblages in phytoplankton, with members of the core microbial lineages (MCGs) playing pivotal roles in shaping host-specific microbial assemblages and contributing to network structures. Deterministic processes, particularly host genotypes, were the dominant factors shaping microbial assemblages, overriding environmental influences. Consequently, microbial composition reflected the evolutionary relationships of the host species, demonstrating phylosymbiotic patterns. These findings suggest that studying MCGs will be a crucial foundation for investigating specific phytoplankton-microbe interactions and highlight the ecological and evolutionary importance of host-microbial specificity in phytoplankton.

## Introduction

Microbial communities in ecosystems play crucial roles in host nutrition, metabolism, protection from pathogens, and regulation of physiological processes such as immunity and development^1, 2, 3, 4, 5^. Host-specific microbial assemblages have been documented in a wide range of organisms, such as vertebrates^6, 7, 8, 9, 10, 11^, invertebrates^9, 12, 13^, and plants^14, 15^, providing clear evidence of host-microbiota coevolution and mutual benefits. Research into host-specific microbiomes is critical for understanding how these microbial partnerships influence the fitness, ecological dynamics, and persistence of host species. In some cases, host specificity is observed as phylosymbiosis, in which microbial assemblages reflect the evolutionary relationships of their hosts^15, 16, 17, 18^. However, this pattern is not universal, as not all marine invertebrates and terrestrial mammals exhibit phylosymbiosis^19, 20^.

As key primary producers in aquatic ecosystems, phytoplankton are responsible for approximately half of Earth’s oxygen production and are crucial for the biogeochemical cycling of carbon, nitrogen, sulphur, and iron^21, 22, 23^. Simultaneously, bacteria play central roles in the decomposition and remineralization of organic matter, forming the functional backbone of these ecosystems^24, 25^. Phytoplankton and bacteria are closely intertwined, with phytoplankton providing dissolved organic carbon, and bacteria supplying essential nutrients such as bioavailable nitrogen and phosphorus, as well as growth-promoting compounds such as vitamins^25, 26, 27^. These reciprocal interactions are particularly evident during algal blooms, during which bacterial communities actively support phytoplankton growth and shape bloom dynamics^27, 28, 29, 30, 31^.

Recent findings have shown that bacteria can promote the growth of specific phytoplankton by regulating their aging physiology, revealing intricate mechanisms underlying these interactions^32^. However, the details of specific interactions between phytoplankton and bacteria remain largely unexplored. A promising starting point for studying these intricate relationships is to examine whether phytoplankton hosts harbour distinct host-specific microbial assemblages.

Although some studies provide evidence for host-specific microbial assemblages in phytoplankton^29, 33, 34, 35, 36, 37, 38^, in part because many analyses were limited to comparisons within a single species or between species using only one strain per species, most research focuses on diatoms, despite the critical ecological roles of dinoflagellates in aquatic systems^39, 40^. Many studies relied on laboratory strains, leaving host-specific microbial assemblages in natural environments largely unexplored.

In this study, we address these gaps by investigating host-specific microbial assemblage patterns in both laboratory-cultured strains and samples of phytoplankton blooms collected from the field. We examined six microalgal species, including diatoms and dinoflagellates, across multiple strains and growth stages, analysing core microbial lineages. Our results provide strong evidence for host-specific microbial assemblages in phytoplankton, even in highly mixed aquatic environments, and identify potential phylosymbiotic patterns between phytoplankton hosts and their microbial assemblages.

## Results

### Host-specificity of microbial assemblages across phytoplankton species

Microbial assemblages associated with six phytoplankton species exhibited distinct compositions depending on the host species (Fig. 1a). Among dinoflagellates, the dominant bacterial taxa were Rhodobacterales (Alphaproteobacteria) and Alteromonadales (Gammaproteobacteria). Sub-dominant taxa varied by species: *Flavobacteriales* (Flavobacteriia) were more abundant in *Alexandrium catenella* (Group I) and *Gymnodinium catenatum*, while members of Cytophagales (Cytophagia) showed higher relative abundance in *A. catenella* and *A. pacificum* (Group IV). In diatoms, Rhodobacterales consistently dominated in *Cylindrotheca closterium* and *Coscinodiscuss granii*, whereas *Alteromonadales* dominated in *Pseudo-nitzschia pungens*. Sub-dominant taxa also varied by species, including Bacillales (Bacilli) in *Cylindrotheca closterium*, Vibrionales (Gammaproteobacteria) in *P. pungens*, and a mix of Alteromonadales and Flavobacteriales in *Coscinodiscuss granii*. Dinoflagellates were significantly enriched in nine bacterial families, including Alteromonadaceae (Gammaproteobacteria), Flavobacteriaceae (Flavobacteriia), and Flammeovirgaceae (Cytophagia), compared with diatoms (linear discriminant analysis [LDA] score > 3.12, adjusted p-value < 0.05). In contrast, Bacillaceae (Bacilli) was the only significantly enriched family in diatoms (Supplementary Fig. 1a). Alpha diversity was significantly higher in dinoflagellates than in diatoms, with dinoflagellates associated with a greater Shannon index (2.54 vs. 1.46) and Chao1 values (125.6 vs. 65.18) (p < 0.001) (Supplementary Fig. 1b and c). This indicates that dinoflagellates harbour a more diverse array of microbial lineages.

**Fig. 1:**
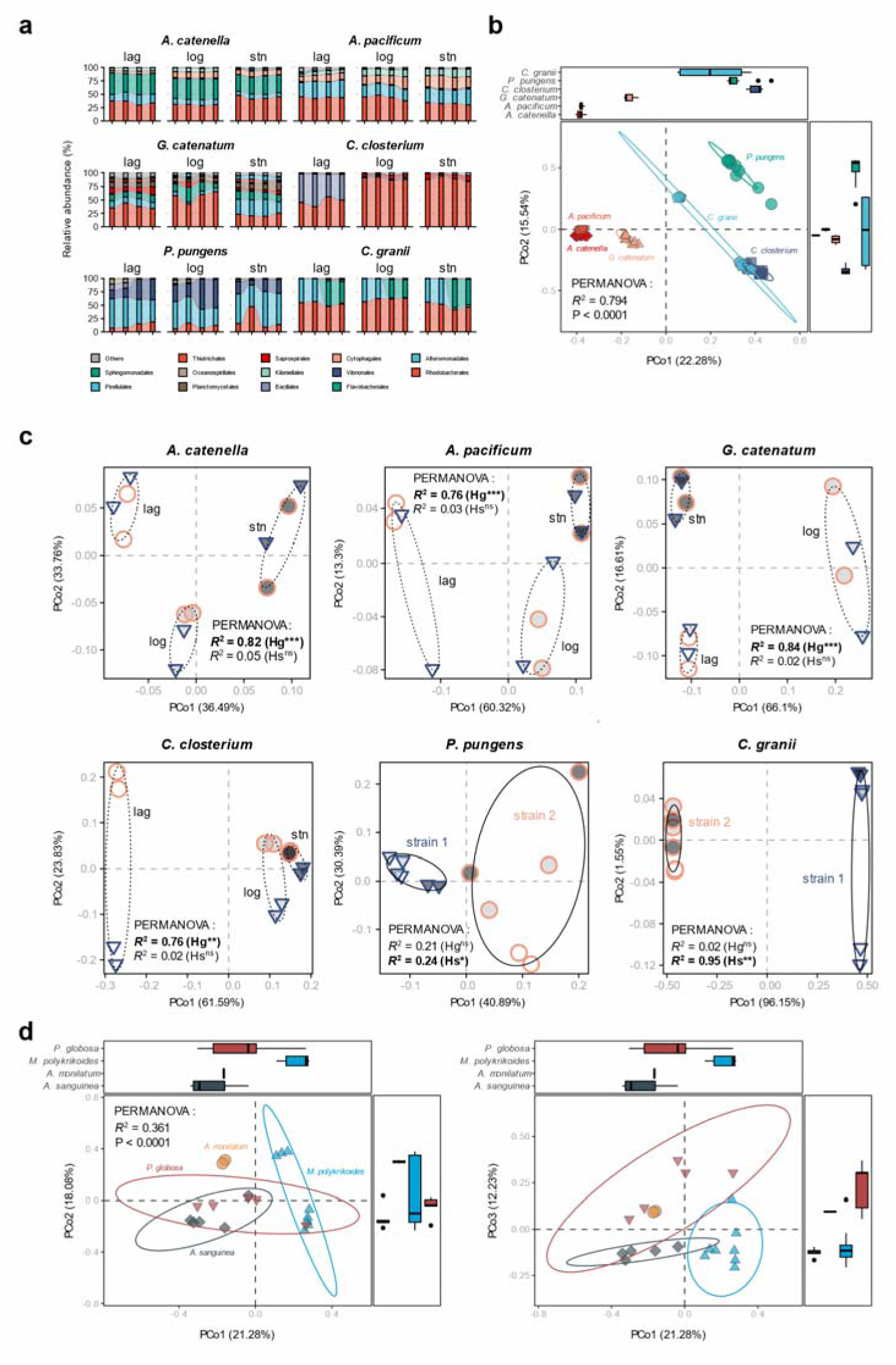
Host-specificity of microbial assemblages in six phytoplankton species, confirmed in both laboratory-cultured strains and field-collected phytoplankton blooms. a: Order-level composition of microbiomes across strains and growth stages of the six phytoplankton hosts. b: PCoA of microbiomes associated with the six phytoplankton species, based on Bray–Curtis dissimilarity. Ellipses represent 95% confidence intervals for each host species. PERMANOVA results indicate significant clustering by host species. c: PCoA highlighting microbiome differences among six phytoplankton species by strain and growth phase. Dashed ellipses indicate growth-phase separation, while solid ellipses represent strain differences. PERMANOVA significance is shown with *, P < 0.05; **, P < 0.01; ***, P < 0.001; ns, not significant. d: PCoA of microbiomes in field-collected blooms of four phytoplankton species (*Akashiwo sanguinea, Alexandrium monilatum*, *Margalefidinium polykrikoides*, *Phaeocystis globosa*) sampled across Korea, China, and the US. Plots display microbiomes along PCoA axes 1 and 2 (left) and axes 1 and 3 (right). Data for samples from China, the US, and some Korean coastal regions were retrieved from the NCBI (Supplementary Table 3). To minimize variability from different sequencing hypervariable regions, microbiome abundance data were collapsed at the genus level.

Beta diversity analysis revealed host-specific microbial assemblages which involve significant differences in the microbial assemblages of each phytoplankton host based on Bray–Curtis dissimilarity and confirmed by principal coordinates analysis (PCoA) and permutational multivariate analysis of variance (PERMANOVA, R² = 0.794, *p* < 0.001) (Fig. 1b). Most microbial assemblages were clustered with the microbial assemblages of the same phytoplankton species isolated from different spatiotemporal contexts (Supplementary Table 1, Supplementary Fig. 2b). For example, although the two strains of *Cylindrotheca Closterium* were separated spatially by 375 km and temporally by 347 days, they were clustered into the same clade in the hierarchical clustering analysis, indicating that they share a conserved microbial assemblage and are distinct from other phytoplankton species (Supplementary Fig. 3). Host-specificity in microbial assemblages was evident not only in Bray–Curtis dissimilarity results but also across other beta diversity metrics, consistently showing statistically significant distinctions (Supplementary Table 2).

Distinct microbial compositions were observed across different growth stages of the host species, highlighting growth-specificity of microbial assemblages. For example, in *A. catenella*, the relative abundance of Rhodobacterales increased, whereas Kiloniellales decreased during the stationary phase compared with the lag phase. This pattern was consistent across all strains and replicates examined (Fig. 1c). PERMANOVA analysis supported these observations, showing that the microbial assemblage during the stationary phase of *A. catenella* strain AC-1 was more similar to the stationary phase microbial assemblage of strain AC-2 than to its own assemblage during the lag or log phases (R² = 0.82, p < 0.001). No significant difference between strains was evident (R² = 0.05, p > 0.05), indicating that the growth stage, rather than strain, plays a dominant role in shaping microbial assemblages. These growth phase–dependent variations were more prominent in dinoflagellates but were only observed in *Cylindrotheca closterium* among diatoms. While the microbiomes of dinoflagellates were highly similar between strains, several strain-specific differences were noted in *P. pungens* and *Coscinodiscuss granii* among diatoms (Fig. 1c). In *P. pungens* and *Coscinodiscuss granii*, no significant differences in microbiome assembly were observed between growth stages, although statistically significant strain-level differences were seen (R² = 0.24, *p* < 0.05; R² = 0.95, *p* < 0.001, respectively).

Host-specific microbial assemblages were evident in phytoplankton bloom samples taken from the field, with microbial communities clustering distinctly according to the host species of the bloom (Fig. 1d, Supplementary Fig. 4). For example, microbial assemblages associated with blooms caused by *Margalefidinium polykrikoides* (found in both Korean coastal waters and those of the United States), *Akashiwo sanguinea* (Korea), *Alexandrium monilatum* (United States), and *Phaeocystis globosa* (China) formed tight clusters distinct from one another in Bray–Curtis dissimilarity-based PCoA analysis (PERMANOVA, R² = 0.36, p < 0.05) (Supplementary Fig. 2, Supplementary Table 3). Microbial assemblages were more strongly differentiated by the host phytoplankton species (R² = 0.36) than by the geographical location of the bloom (R² = 0.27), highlighting the dominant influence of host species on community structure (p < 0.05).

### Core and host-specific microbial lineages across dinoflagellates and diatoms

In laboratory-cultured strains, the number of core microbial lineages (core MLs) varied by host species, ranging from 5 to 33, excluding *Coscinodiscuss granii*, which lacked observable core MLs (Fig. 2a). Host-specific microbial lineages (HS-MLs) which were statistically significantly different and uniquely associated with individual hosts ranged from 1 to 16 (Fig. 2a, 2b). Notably, nearly all HS-MLs (45 of 47 microbial lineages) overlapped with core MLs, highlighting their strong association (Fig. 2a). Among the top 200 microbial lineages for each host species, the proportion of core MLs and HS-MLs, collectively termed the members of the core MLs group (MCG), ranged from 3% to 17%, indicating that MCGs were numerically fewer than non-MCGs (Fig. 2c). However, MCGs accounted for a substantial portion of the total rarefied abundance for each host species, contributing 80.8% to 96.2%, suggesting that MCGs, while limited in diversity, dominate the microbial community quantitatively.

**Fig. 2:**
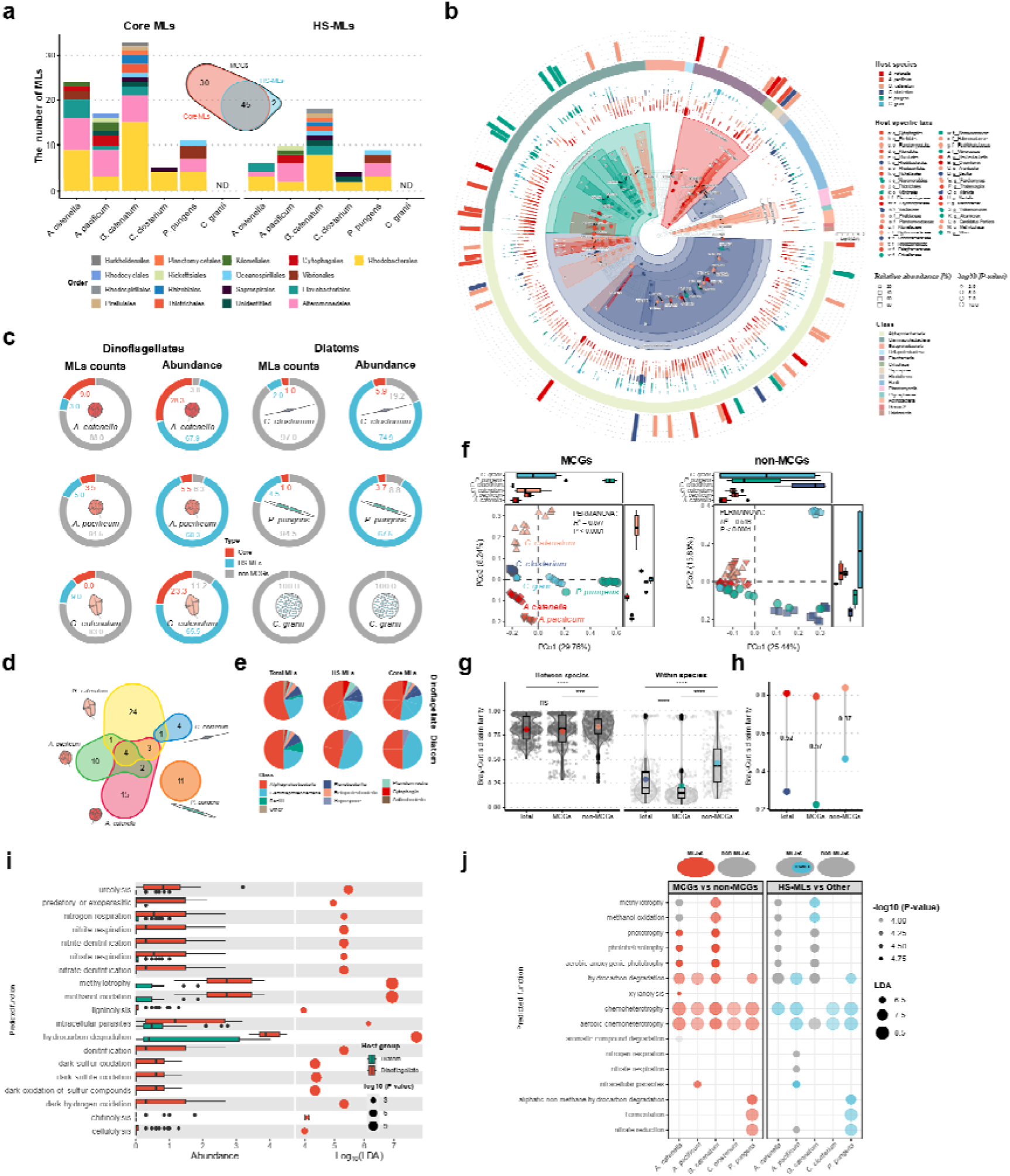
Members of core microbial lineages (MCGs) play a key role in shaping host-specific microbial assemblages across phytoplankton species. a: Number of core MLs and HS-MLs at the bacterial order level for each host species. ND indicates not detected. The central Venn diagram shows the overlap between Core MLs and HS-MLs. **b:** Microbiome taxa tree highlighting host-specific taxa. The outermost points of the tree represent the relative abundance of each microbial lineage. Nodes are colour-coded to indicate statistically enriched taxa for each host species, with node size reflecting the adjusted p-value (Bonferroni correction). The external bar plot shows linear discriminant analysis scores of differentially abundant taxa, with an effect size threshold of 3.2. Differential abundance analysis was conducted using a two-step approach: a Kruskal–Wallis test to detect taxa with overall group differences, followed by a Wilcoxon rank-sum test for pairwise comparisons. **c**: Proportion of core MLs and HS-MLs among the top 200 MLs for each host species by number of MLs and relative abundance. Core MLs shown here exclude HS-MLs within the MCGs. **d:** Venn diagrams of Core MLs for each host species. Circle sizes are proportional to the numbers indicated. *Coscinodiscuss granii*, which lacks core MLs, is not represented. **e:** Bacterial composition by lineage group (ML types) within dinoflagellates and diatoms. Percentages were calculated based on the number of microbial lineages. **f:** Bray– Curtis dissimilarity-based PCoA plots of MCGs (left) and non-MCGs (right) datasets. PERMANOVA results indicate significant clustering by host species. **g:** Distribution of Bray–Curtis dissimilarity values between and within species for the total microbiome, MCGs, and non-MCGs datasets. Boxplot boundaries represent the first and third quartiles, with the median indicated. Whiskers extend to the furthest data points within 1.5× the interquartile range. Coloured points inside the boxes denote mean values (each calculated from *n*=630). **h:** Comparison of between-species and within-species Bray–Curtis dissimilarity values for the total microbiome, MCGs, and non-MCGs datasets. The endpoints of the dumbbells represent the mean values from panel g. The values at the midpoints of the dumbbells represent the differences between between-species and within-species dissimilarities. **i:** Bacterial functional group abundances across phytoplankton species, as predicted by FAPROTAX analysis. Of the 19 functional groups that differed significantly between dinoflagellates and diatoms, all were more abundant in dinoflagellates on average. The left panel depicts the differences in functional group abundance, while the right panel presents linear discriminant analysis scores alongside adjusted p-values (Benjamini-Hochberg correction). Differential abundance was determined using a two-step approach: first, a Kruskal–Wallis test to identify group-level differences, followed by pairwise comparisons using a Wilcoxon rank-sum test. Boxplot boundaries represent the first and third quartiles, with the median indicated, and whiskers extend to the furthest data points within 1.5× the interquartile range. **j:** Differential abundance analysis of bacterial functional groups between ML groups (left: MCGs vs. non-MCGs; right: HS-MLs vs. other) using FAPROTAX predictions. Bubble colours indicate significantly enriched functional groups (adjusted p-value ≤ 0.05). To reduce visual complexity, nonsignificant results (p-value > 0.05) and groups with small effect sizes (LDA score < 6.28) are not shown.

MCGs shared between different hosts were also observed. The number of MCGs shared between host pairs ranged from 1 to 3, with nearly all shared MCGs found exclusively within dinoflagellates (Fig. 2d). Additionally, four MCGs were identified in the dinoflagellates *A. catenella*, *A. pacificum*, and *G. catenatum*, three of which belonged to the family Alteromonadaceae, genus *Marinobacter* (the fourth was from an unidentified genus within Alteromonadaceae). In contrast, no shared MCGs were found among diatom hosts. Dinoflagellates consistently harboured more MCGs (17–34 per species) compared with diatoms (0–11 per species).

Among dinoflagellates, the bacterial taxa associated with MCGs primarily included Rhodobacterales (Alphaproteobacteria), Alteromonadales (Gammaproteobacteria), and Flavobacteriales (Flavobacteriia), closely reflecting the taxonomic composition of all the microbial lineages detected in dinoflagellates (Fig. 2e). In contrast, diatoms exhibited a more restricted and simplified taxonomic composition within their MCGs. For example, while Bacilli and Flavobacteriia constituted 9.1% and 8.5%, respectively, of the total microbial lineages observed in diatoms, neither was represented among MCGs. Instead, MCGs in diatoms were dominated by Rhodobacterales (50.0%) and Alteromonadales (43.8%), suggesting that the core microbiome of diatoms is not only smaller but more taxonomically constrained compared with that of dinoflagellates.

MCGs played a major role in driving host-specific microbial assemblages in phytoplankton species. Although MCGs collectively accounted for only 77 of the top 242 MLs observed in this study, a PCoA and PERMANOVA based solely on these MLs found stronger differentiation among host species compared with analyses of the remaining 165 MLs (non-MCGs) (Fig. 2f). Specifically, weighted UniFrac-based PERMANOVA results for MCG (R² = 0.877, *p* < 0.0001) showed significantly greater intergroup differences compared with those observed for the remaining MLs (R² = 0.516, *p* < 0.0001) or the full dataset of 242 MLs (R² = 0.794, *p* < 0.001). Consistent trends were also observed across beta diversity metrics, such as Bray–Curtis dissimilarity, further supporting the robustness of these findings (Table 1). The difference in Bray–Curtis dissimilarity between species and within species was larger when calculated using MCG compared with calculations using non-MCGs or the total microbiome, highlighting the high contribution of MCGs to host specificity (Fig. 2g and h).

**Table 1.** PERMANOVA results comparing microbial community structure across groups and distance metrics.

| Group | Number of MLs | Distance metrics | Pseudo F | R<sup>2</sup> | P-value |
| --- | --- | --- | --- | --- | --- |
| Total MLs | 242 | Bray-Curtis | 50.884 | 0.794 | p < 0.0001 |
| Total MLs | 242 | Weighted UniFrac | 49.862 | 0.791 | p < 0.0001 |
| Total MLs | 242 | Unifrac | 22.225 | 0.627 | p < 0.0001 |
| MCGs | 77 | Bray-Curtis | 61.228 | 0.823 | p < 0.0001 |
| MCGs | 77 | Weighted UniFrac | 93.756 | 0.877 | p < 0.0001 |
| MCGs | 77 | UniFrac | 21.098 | 0.615 | p < 0.0001 |
| non-MCGs | 165 | Bray-Curtis | 18.053 | 0.578 | p < 0.0001 |
| non-MCGs | 165 | Weighted UniFrac | 14.088 | 0.516 | p < 0.0001 |
| non-MCGs | 165 | UniFrac | 28.860 | 0.686 | p < 0.0001 |

### Functional group differences between dinoflagellate and diatom microbiomes and between core MLs group and other MLs

Functional annotation of prokaryotic taxa (FAPROTAX) analysis revealed significant differences in bacterial functional groups between dinoflagellates and diatoms (Fig. 2i, Supplementary Fig. 5). Notably, all functional groups showing significant differences were more enriched in dinoflagellates compared with diatoms. The functional group with the greatest difference between these two microalgal groups, based on LDA effect size, was “hydrocarbon degradation”. Other notable functional groups enriched in dinoflagellates included those related to carbon metabolism, such as “methylotrophy” and “methanol oxidation”, as well as groups associated with nitrogen and sulphur recycling and biopolymer degradation.

Core MLs exhibited distinct functional profiles compared with other MLs, with chemoheterotrophic functions being significantly enriched across all host species (Fig. 2j). Additionally, hydrocarbon degradation was a prominent function observed in the core MLs of most hosts. Some functional differences among core MLs were also noted across hosts. For example, core MLs in *A. catenella* were enriched in “xylanolysis”, while those in *A. pacificum* exhibited functions associated with “intracellular parasites“. In *G. catenatum*, “methylotrophy“-related functions were prominent, whereas core MLs in *P. pungens* were enriched in functions related to “fermentation” and “nitrate reduction“. Functions observed in HS-MLs overlapped significantly with those in core MLs, including key functions enriched in specific hosts, such as hydrocarbon degradation and chemoheterotrophy.

### Network analysis of microbiomes by microalgal host species

Co-occurrence network analysis of the microbiomes associated with each microalgal host revealed that bacterial network connectivity within dinoflagellate microbiomes was generally weaker compared with diatom microbiomes (Fig. 3a). Specifically, the clustering coefficient was significantly lower in dinoflagellates (0.57) than in diatoms (0.73) (Fig. 3b). Although the average degree (31.68 in dinoflagellates versus 65.46 in diatoms) and connectance (0.16 in dinoflagellates versus 0.33 in diatoms) also differed, these differences were not statistically significant.

**Fig. 3:**
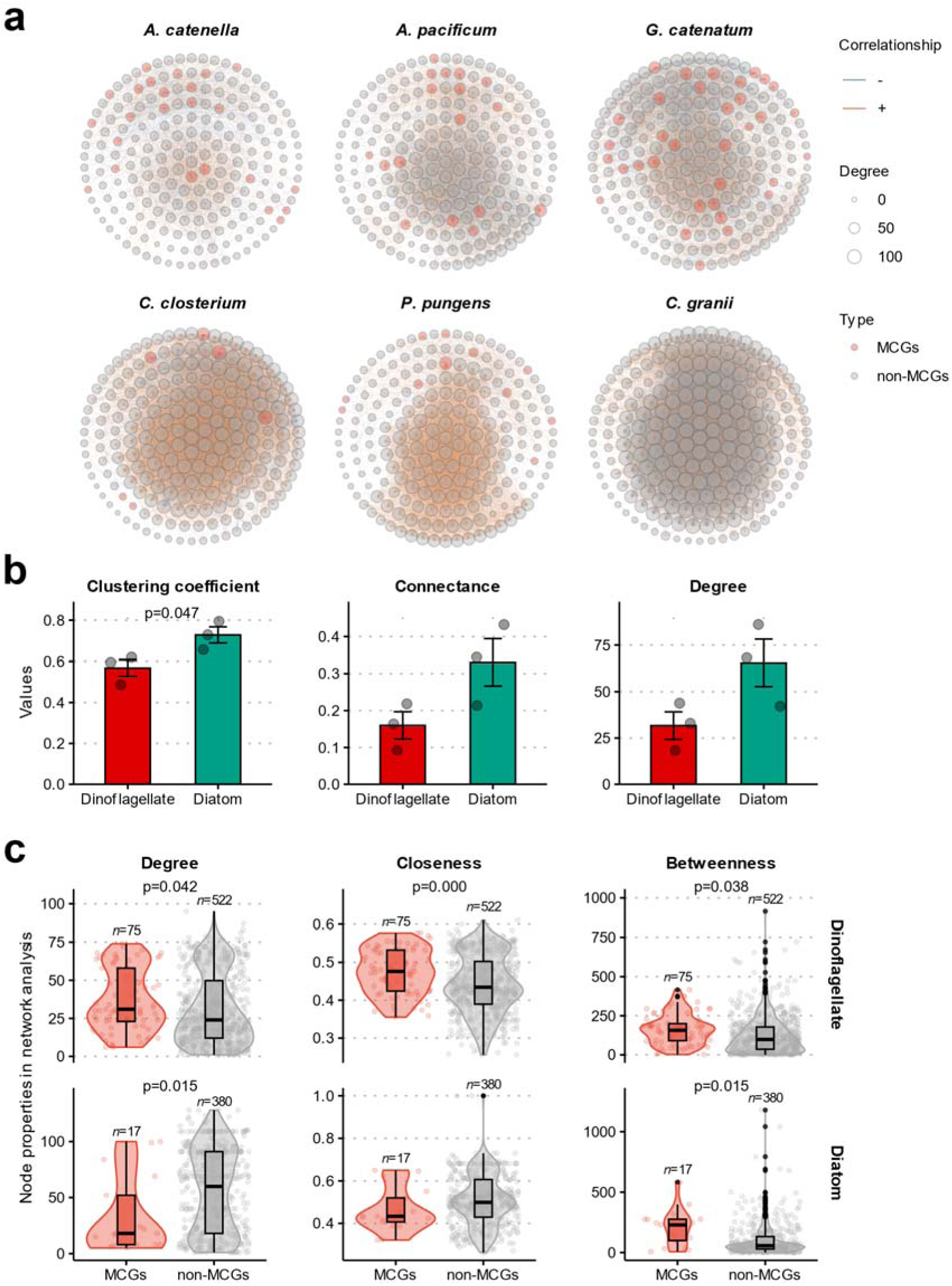
Properties of co-occurrence networks and the significance of MCG nodes in phytoplankton-associated microbiomes. a: Co-occurrence networks constructed from the top 200 microbial lineages by abundance for each of the six phytoplankton hosts. The upper row represents microbiomes associated with dinoflagellates, while the lower row depicts those of diatoms. Nodes are colour-coded to distinguish MCGs from non-MCGs lineages. *Coscinodiscuss granii* contains only non-MCG nodes as no MCGs were identified. **b:** Comparison of three network properties (clustering coefficient, connectance, and degree) between dinoflagellate-and diatom-associated microbiomes. Data represent the mean values for each group (n = 3). Error bars indicate standard errors. Numbers above the bars denote p-values. Statistical significance was calculated using a two-sided Student’s t-test. **c:** Distribution of three node properties (degree, closeness, and betweenness) for MCG and non-MCG nodes within each host’s co-occurrence network. Boxplot boundaries represent the first and third quartiles, with the median indicated. Whiskers extend to the furthest data points within 1.5× the interquartile range. To evaluate statistical significance and address differences in sample sizes between groups, a permutation test with 1,000 resamples was performed. P-values were adjusted using the Benjamini–Hochberg method.

Within dinoflagellate microbiome networks, MCG nodes exhibited significantly higher degree, closeness, and betweenness centrality compared with other nodes, emphasizing their critical role in network structure (Fig. 3c). In contrast, in diatom microbiomes, other nodes showed higher degrees and closeness centrality compared with MCG nodes, indicating that the relative importance of MCG nodes in diatom networks was lower than in dinoflagellates.

### Deterministic influences and phylosymbiotic patterns in microalgal microbiome assembly

To evaluate the relative contributions of stochastic and deterministic processes in shaping the host-specific microbiome assembly, we analysed the normalized stochasticity ratio (NST) of the Jaccard matrix (NST_jac_). This metric distinguishes between stochastic (NST > 50%) and deterministic (NST < 50%) influences on community structure. The results indicate that microbiome structures in both dinoflagellates and diatoms were predominantly shaped by deterministic factors (Fig. 4a). Dinoflagellates exhibited a mean NST value of 16.8 ± 3.1%, with a maximum value of 32.3%, reflecting a strong deterministic control. In contrast, diatoms showed a higher mean NST value of 32.4 ± 13.2% and a maximum of 90.1%, suggesting that stochastic influences were more prominent in diatom microbiomes compared with dinoflagellates.

**Fig. 4:**
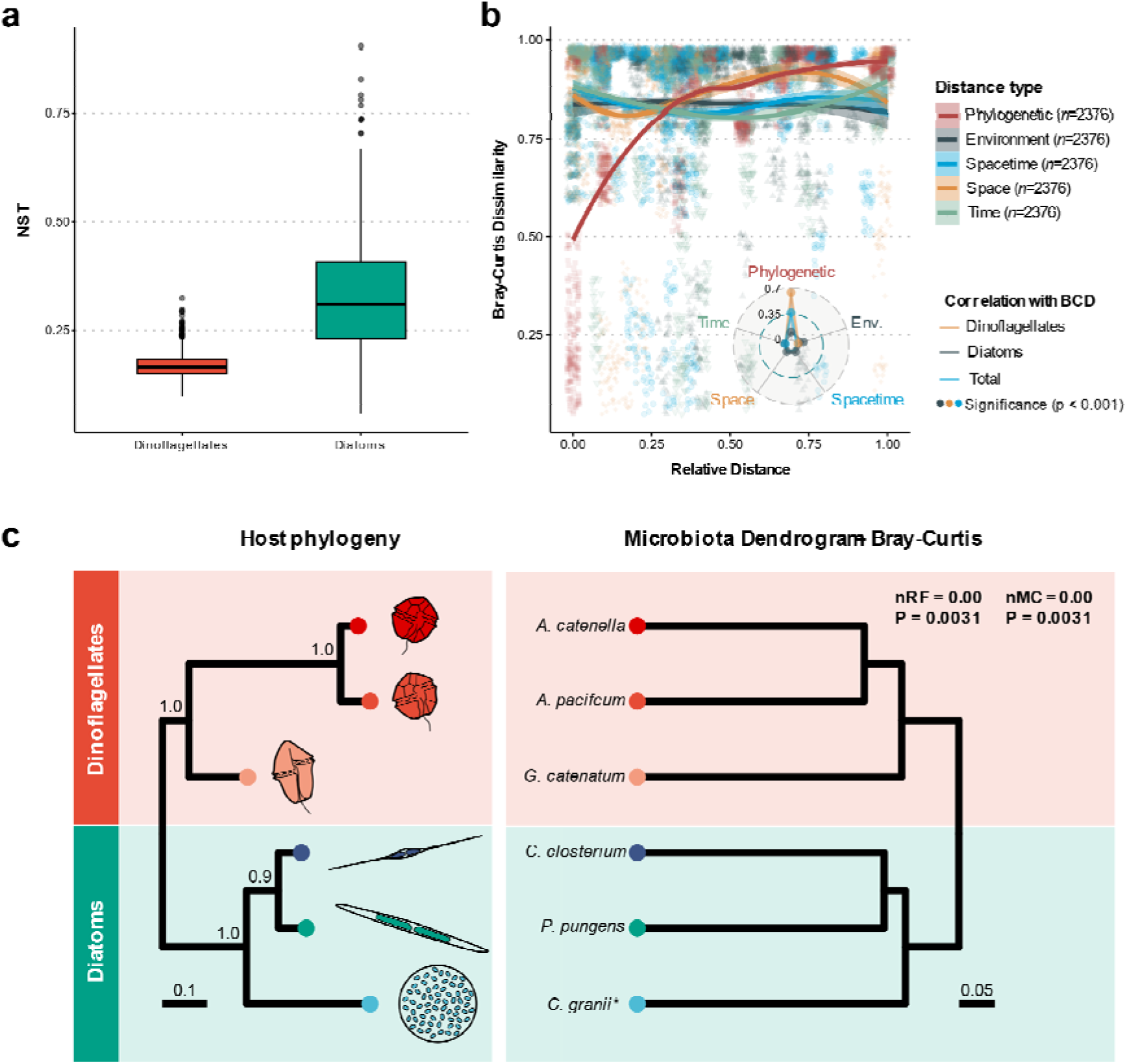
Microbiome distance in phytoplankton is most strongly correlated with the evolutionary distance between hosts, supporting the observation of phylosymbiotic patterns. **a**: Comparison of the normalized stochasticity ratio (NST) of the Jaccard matrix between dinoflagellates and diatoms. Values greater than 0.5 indicate that stochastic factors play a dominant role in microbiome assembly, while values below 0.5 indicate a stronger influence of deterministic factors. **b:** Correlation of Bray–Curtis dissimilarity between host microbiomes with phylogenetic, physicochemical, and spatiotemporal distances. The phylogenetic distance between hosts was calculated using pairwise distances based on LSU rDNA sequences (1,031 bp). All distance types, except for Bray–Curtis dissimilarity and phylogenetic distance, were measured as Euclidean distances and normalized to a maximum value of 1. The R² values for each distance type’s correlation with Bray–Curtis dissimilarity are displayed in radar plots, categorized by dinoflagellates, diatoms, and total data. All significant correlations had P < 0.001. Lines show loess regression fits between Bray–Curtis dissimilarity and each type of distance, with shaded areas represent the 95% confidence intervals. **c:** Phylosymbiotic analysis comparing topological congruence between the rooted host phylogeny and microbiota dendrogram. The host phylogenetic tree was constructed using LSU rDNA sequences (1,031 bp), and the microbiota dendrogram was generated by UPGMA clustering based on Bray–Curtis dissimilarity. Topological congruences were quantified using the normalized Robinson–Foulds (nRF) and normalized matching cluster (nMC) metrics, with values ranging from 0 (complete congruence) to 1 (complete incongruence). *Note: LSU rDNA sequence data for* Coscinodiscuss granii *was unavailable in NCBI, so the sequence from its closely related species,* Coscinodiscuss wailesii*, was used instead*.

Correlation analysis to identify deterministic factors influencing microbial assemblages revealed that the dissimilarity between microalgal-associated microbiomes was not significantly correlated with physicochemical factors, including temperature, salinity, pH, dissolved oxygen (DO), total nitrogen (TN), total phosphorus (TP), chemical oxygen demand (COD), chlorophyll-a (Chl-a), or spatiotemporal distance at the time of host strain isolation. However, a strong positive correlation was observed with the phylogenetic distance between hosts (R = 0.61, *p* < 0.001) (Fig. 4b). Specifically, microbiome similarity increased with decreasing evolutionary distance between hosts. This relationship was notably stronger in dinoflagellates (R = 0.80, *p* < 0.001) compared with diatoms (R = 0.33, *p* < 0.001), indicating a more pronounced influence from host genetic relatedness on microbiome assembly in dinoflagellates.

To further investigate phylosymbiotic patterns, we performed Robinson–Foulds and matching cluster analyses, which showed complete congruence (normalized distance = 0.0) between the molecular phylogeny of the host phytoplankton and the dendrogram topology of their microbiome dissimilarity across both evaluation methods (nRF = 0.00, p < 0.01; nMC = 0.00, p < 0.01) (Fig. 4c). The microbiome dendrograms could be divided into two major groups: dinoflagellates and diatoms. Within these groups, the microbiomes reflected the phylogenetic traits of their hosts, with dinoflagellate microbiomes clustering into unarmoured and armoured subgroups, while diatom microbiomes separated into pennate and centric subgroups. Additionally, congruence was also observed when using unweighted UniFrac to calculate microbiota distances, although no significant congruence was detected with weighted UniFrac (Table 2).

**Table 2.** The normalised Robinson–Foulds (nRF) and the normalized Matching Cluster metric analysis across Bray–Curtis, weighted and unweighted UniFrac beta-ersity metrics.

| Metric | Bray-Curtis | Weighted UniFrac | Unweighted UniFrac |
| --- | --- | --- | --- |
| nRF | RF = 0.00, p = 0.0031 | RF = 0.50, p = 0.0735 | RF = 0.00, p = 0.0031 |
| nMC | MC = 0.00, p = 0.0031 | MC = 0.83, p = 0.9637 | MC = 0.00, p = 0.0031 |

## Discussion

In this study, through comparative analyses of both laboratory-cultured strains and field-collected bloom samples, we found evidence suggesting that microbial assemblage in phytoplankton exhibited a degree of host-specificity. We found that strains of the same microalgal species, isolated from spatiotemporally distant locations, exhibited highly similar microbiome assemblies. Even phytoplankton strains isolated from the same time and location, such as *A. catenella* (AC-2), *A. pacificum* (AP-2), and *G. catenatum* (GC-1), shared only 4 MLs, and these MLs were consistent detected in MLs of strains isolated from different times and locations, such as *A. catenella* (AC-1), *A. pacificum* (AP-1), and *G. catenatum* (GC-2), indicating microbial assemblages in phytoplankton are highly preserved and have their own specificity (Supplementary Fig. 6). The region surrounding phytoplankton cells, known as the phycosphere, is rich in organic matter, which is known to support active interactions with bacteria^27, 41^. Our findings suggest that the phytoplankton phycosphere is remarkably robust in maintaining its microbial assemblages, likely facilitated by active host-microbe interactions within this microenvironment.

While previous studies have explored host-specific microbial assemblages in phytoplankton, they primarily focused on single species or limited geographical regions^28, 29, 36, 38, 42, 43, 44^. By analysing samples of bloom samples collected from multiple geographically distinct locations in the field, our study provides novel insights into host-specificity across multiple phytoplankton species under natural conditions. Notably, while the similarity of microbial assemblages within the same species was moderately lower in samples of phytoplankton blooms compared with lab-cultured strains, the degree of similarity remained high in natural environments. This difference can be attributed to the methodologies used for sample preparation. Lab-cultured strains, established through single-cell isolation, are expected to retain microbes that are physically closely associated with the host cells. In contrast, field-collected bloom samples, analysed by filtering entire seawater samples, likely included microbes unrelated to the host. As a result, host specificity in microbial assemblages was moderately reduced in field samples. Despite this reduction, microbial assemblages associated with the same phytoplankton species exhibited remarkable similarity across field samples, even when collected from geographically and temporally distant locations. This underscores the robustness of host-specific microbial assemblages in phytoplankton, even under the dynamic conditions of natural environments.

The observation that phytoplankton, which inhabit a homogenous water column, maintain host-specific microbial assemblages is particularly intriguing. Most studies reporting host-specific microbial assemblages focused on heterogeneous environments, such as the gut microbiomes of vertebrates and invertebrates, or root-associated microbiomes of terrestrial plants^6, 13, 14, 15, 18, 45, 46, 47^. Our findings indicate that even phytoplankton, which lack organs such as a gut, can exert a significant influence on the structure of their associated microbiomes.

A key factor underlying the host specificity observed in phytoplankton microbiomes is the role of MCGs, which dominate host-associated microbial assemblages and contribute significantly to their distinct composition. Despite representing a smaller fraction of microbial diversity compared with non-MCGs, MCGs dominate host microbiomes and contribute significantly to their host specificity. The majority of MCGs identified in our study belong to the orders Rhodobacterales, Alteromonadales, and Flavobacteriales, which have been frequently associated with phytoplankton cultures and blooms in previous studies^25, 48, 49, 50^. Among these, members of the family Roseobacteraceae (Order Rhodobacterales) dominated the Alphaproteobacteria class, accounting for 86 of 122 microbial lineages. Roseobacteraceae species are known to utilize various organic compounds released by phytoplankton, such as dimethylsulphoniopropionate and amino acids, and to participate in key biogeochemical processes involving carbon, nitrogen, sulphur, and phosphorus cycling. Additionally, they produce vitamins and growth-promoting compounds that benefit phytoplankton^51, 52^. Alteromonadaceae (Order Alteromonadales) break down diverse polysaccharides, using carbohydrate-active enzymes encoded in polysaccharide utilization loci^53^. This group is particularly adept at degrading complex biopolymers, such as laminarin, alginate, xylan, and mannans, facilitated by their large genomes and extensive enzymatic repertoire. Similarly, Flavobacteriia use polysaccharides and proteins as primary sources of carbon and energy, efficiently recycling organic material and nutrients from phytoplankton blooms by processing photosynthetic byproducts or degrading algal cell walls^30, 54, 55, 56^. Together, these MCGs are likely to play complementary roles in the remineralization of organic matter within the phytoplankton microenvironment, potentially contributing to localized nutrient cycling and energy flow.

Beyond the dominance of MCGs in host microbiomes, their ecological significance lies in their diverse functional capabilities, which can be characterized using FAPROTAX predictions and reveal key metabolic traits crucial for interactions with phytoplankton hosts. Across all host species, MCGs exhibited distinct functional profiles compared with non-MCGs, with chemoheterotrophic functions being significantly enriched. This indicates that MCGs are likely highly dependent on host-derived organic matter, and suggests a commensal or even mutualistic relationship between phytoplankton hosts and their associated MCGs. Hydrocarbon degradation was a prominent function consistently observed in the MCGs of most hosts, far exceeding its representation in non-MCGs. Among dinoflagellates, the genus *Marinobacter* (family Alteromonadaceae) was commonly detected in the MCGs of all host species. *Marinobacter* is a well-known hydrocarbon-degrading bacterium, and one lineage, OTU22, was found to share 99.57% of its gene sequence with *Marinobacter algicola*, a hydrocarbon-degrading bacterium previously isolated from *Alexandrium tamarense* and *Gymnodinium catenatum*^57^. This highlights the inclusion of hydrocarbon-degrading bacteria in the core microbiome of dinoflagellates, a result consistent with previous studies^42, 58^. Such associations suggest a particularly close ecological relationship between these microbial lineages and dinoflagellates. Some phytoplankton species, including dinoflagellates, are known to produce hydrocarbons^59, 60, 61, 62^, which may explain the frequent presence of bacteria capable of hydrocarbon degradation. However, it remains unclear whether such bacteria are associated with all phytoplankton because they metabolize host-derived hydrocarbons or because they provide essential benefits critical for host survival. Previous studies have shown that hydrocarbon-degrading bacteria can promote phytoplankton growth by supplying essential nutrients or growth-promoting compounds^63, 64^. Further research is needed to clarify whether these interactions constitute a mutualistic relationship and elucidate the underlying mechanisms driving these associations.

As revealed by a microbiome co-occurrence network analysis, the MCG nodes in dinoflagellates played more significant roles in network connectivity compared with other nodes. This suggests that MCGs may act as ecological bridges, not only mediating interactions with the host but facilitating connections among other bacterial taxa. MCGs appeared to contribute substantially to maintaining the overall microbial assemblages of their hosts. Meanwhile, the co-occurrence network results for dinoflagellates, in which bacterial networks were weaker than those observed in diatoms, imply that dinoflagellate-associated bacterial communities are more tightly regulated by the host. This pattern resembles the weakening of microbial network stability seen under environmental stress, in which microbial assemblages across samples exhibit minimal variation and high similarity ^65^.

The weaker network connectivity observed in dinoflagellate microbiomes can be attributed to their host-specific microbial assemblages, which are more pronounced compared with those in diatoms. This is supported by the larger number of MCGs found in dinoflagellates and the higher similarity values within species than between species. The within-species similarity observed in our study aligns with that reported previously in research on microalgal microbiomes. The bacterial community structures associated with dinoflagellates were largely consistent at the order level, while diatoms exhibited greater variability. For example, in *A. catenella*, both our study and previous research identified Rhodobacterales, Cytophagales, and Flavobacteriales as dominant taxa^44^. In contrast, during the exponential phase of *Cylindrotheca closterium*, Rhodobacterales consistently emerged as the dominant taxon, aligning with our findings. However, unlike *A. catenella*, the composition of secondary dominant taxa in *Cylindrotheca closterium* varied between studies^43^. In *P. pungens*, earlier studies reported Rhodobacterales was the dominant taxon, with minimal representation of Alteromonadales. In our study, however, Alteromonadales was dominant, with Rhodobacterales being subdominant^33^. This suggests that diatom microbiomes are less conserved and more dynamic compared with dinoflagellate microbiomes.

These findings align with previous reports indicating that diatom microbiomes, while generally conserved, can vary depending on environmental conditions at the time of strain isolation^43, 66^. This flexibility in microbiome composition was most pronounced in *Coscinodiscuss granii*, which lacked observable core MLs, whereas the microbiomes of the other two diatom species were more stable. This observation does not entirely align with studies reporting a highly conserved microbiome structure in diatoms^33, 34, 35^. However, the taxonomic diversity within diatoms, categorized into three major classes — Coscinodiscophyceae, Mediophyceae, and Bacillariophyceae — provides a plausible explanation^67, 68^. In this study, *Coscinodiscuss granii* belongs to Coscinodiscophyceae, while the other two species are members of Bacillariophyceae. The variation in microbiome conservation observed across diatom species and studies may reflect evolutionary and coevolutionary dynamics between hosts and their microbiomes, underscoring the need for further research to elucidate these relationships^47, 69, 70^.

The reason for the greater microbial assemblage similarity among dinoflagellate hosts compared to diatom hosts is uncertain. Many animals and plants regulate their microbiomes to benefit the host by producing metabolites, such as adhesive molecules, antimicrobial peptides, and hormones^15, 71, 72, 73^. Similarly, host-specific microbial assemblages in phytoplankton are thought to be strongly shaped by host-derived metabolites. For example, diatoms selectively modulate surrounding bacteria through the production of secondary metabolites^74^. Other studies demonstrated that manipulating the ratios of representative metabolites produced by two phytoplankton species can not only alter the bacterial community composition in natural seawater consortia but also produce predictable outcomes^75, 76^. This serves as an example of how microbial assemblages can be regulated by phytoplankton hosts.

The relatively strong host specificity observed in the microbiomes of dinoflagellates compared with that in diatoms may suggest that dinoflagellate-derived metabolites exert stronger selective pressures and that dinoflagellates may rely more heavily on their microbiomes. Results of a FAPROTAX analysis showed that dinoflagellates harbour a much richer array of bacterial functional groups compared with diatoms. Although the specific benefits these functional groups provide to the host remain unclear, dinoflagellates may depend more heavily on their microbiome’s functions and may apply stricter filtering to recruit beneficial bacteria^77^.

The microbial assemblage is influenced by various factors, including selection, dispersal, diversification, and drift ^78, 79, 80, 81^. These factors are typically categorized into deterministic and stochastic processes. While traditional niche-based ecological perspectives emphasize that environmental factors play a significant role in shaping communities through deterministic processes^82^, recent studies suggest that stochastic factors may better predict variations in community structure^83, 84, 85^. In this study, microbial assemblages observed in both dinoflagellate and diatom microbiomes were predominantly governed by deterministic processes. However, NST analysis of diatom microbiomes revealed a greater prevalence of stochastic influences, with values exceeding the 50% threshold. This suggests that diatom microbiomes may be shaped by priority effects and other stochastic factors^38, 86^.

In our analysis, deterministic environmental factors such as temperature, salinity, pH, DO, COD, TN, TP, and Chl-*a*, as well as the spatiotemporal location of strain isolation, were considered. However, the factor most strongly correlated with microbiome dissimilarity (Bray–Curtis) between samples was not the physicochemical environment or spatiotemporal distance, but the evolutionary distance between hosts. This aligns with the results of previous studies showing that a host’s genotype can significantly influence its microbiome^38^. It also suggests that microalgal microbiomes are shaped primarily by host filtering, with selection acting as the dominant deterministic factor in microbiome assembly.

The proposition that microbiome similarity increases as evolutionary relationships between hosts converge is central to the emerging idea of phylosymbiosis, which has attracted significant attention from ecologists over the past decade^70, 86, 87^. Phylosymbiosis has been observed in various organisms, including insects, mammals, and marine invertebrates, although some taxa do not exhibit this pattern ^9, 20, 88^. To our knowledge, this is the first study to suggest that phytoplankton associated microbiomes may exhibit phylosymbiotic characteristics. We found evidence of phylosymbiosis in phytoplankton microbiomes despite their unicellular structure and microscopic size. This contrasts with findings reported for marine microscopic invertebrates, where such features are absent^20^. However, the congruence between microbiota dendrograms and host phylogenies varied depending on the beta diversity metrics used, highlighting the need for further validation. Given that this study examined only six host species and two strains per species, additional research with a broader range of host taxa and multiple strains is required to confirm the phylosymbiotic nature of phytoplankton-associated microbiomes. Moreover, future studies should explore whether phylosymbiosis is a universal trait among phytoplankton and identify the mechanisms driving this pattern, such as host-derived metabolites or ecological interactions shaping microbial community assembly.

## Conclusion

This study identifies MCGs in phytoplankton microbiomes and describes their contribution to host-specific microbial assemblages. Investigating the interactions between host-specific MCGs and their phytoplankton hosts will be critical for elucidating previously unknown mechanisms of phytoplankton-bacteria interactions. Additionally, phylosymbiotic patterns were observed between phytoplankton and their microbiomes, suggesting a more interdependent co-evolutionary relationship than previously recognized. These findings underscore the necessity of jointly considering phytoplankton and bacterial communities to comprehensively understand their ecological dynamics and evolutionary interplay in marine environments.

## Methods

### Microalgal strain collection and bloom sampling

To analyse the microbiomes of phytoplankton, three dinoflagellate species and three diatom species were selected (hereafter referred to as “lab microbiomes”). To minimize the impact of environmental factors on microbiome similarity, two strains with different spatiotemporal origins were used for each microalgal host species. For example, two strains of *Cylindrotheca closterium* were isolated from locations 375 km and 347 days apart. All algal strains were obtained from the Library of Marine Samples of the Korea Institute of Ocean Science and Technology in Geoje, South Korea. Information on the environmental conditions at the time of strain isolation (e.g., temperature, salinity, pH, dissolved oxygen, COD, TN, TP, and Chl-a) was provided (Supplementary Table 1). All strains were cultured in duplicate at 20°C in an F/2 growth medium ^89^ with a salinity of 31–33 under cool-white fluorescent lamps (photon flux of 100 μE m^−2^ s^−1^) on a 12-h light/12-h dark cycle.

To observe microbiome composition changes across growth stages for each host, samples of 300 mL were taken during the lag, log, and stationary phases based on cell density. The samples were size-fractioned using 47 mm diameter Isopore membrane filters (Millipore, Cork, Ireland) with a pore size of 0.22–3.0 µm, and the filters were stored in an EX buffer (100 mM Tris-HCl, 100 mM Na2-EDTA, 100 mM sodium phosphate, 1.5 M NaCl, 1% CTAB) at −80°C until DNA extraction^90^.

To compare microbiomes across field samples (“field microbiomes”), blooms of *M. polykrikoides*, a major harmful algal bloom dinoflagellate species in Northeast Asia, were collected from four different locations along the Korean coast in 2012 and 2015 (Supplementary Table 4). Additionally, samples of *Akashiwo sanguinea* (Dinophyceae) blooms from 2011 were included for comparison. Seawater samples were collected at a depth of <1 m using a 4.2 L Van Dorn water sampler (Wildlife Supply Company, MI, USA). Volumes of 50–300 mL of seawater were filtered by gravity onto Isopore filters (Millipore, Cork, Ireland) with a pore size of 5 µm, and the filters were submerged in the EX buffer and stored at −80°C until further analysis.

In addition to the field samples, previously published microbiome data from other microalgal blooms were included for comparison. Microbiome data from *M. polykrikoides*, *Alexandrium monilatum*, *A. sanguinea,* and *Phaeocystis globosa* (Prymnesiophyceae) blooms were retrieved from the National Center for Biotechnology Information (NCBI) Sequence Read Archive (PRJNA731462, PRJNA1061360, PRJNA685631, PRJNA678802)^28, 91, 92, 93^ (Supplementary Table 3).

### DNA extraction and sequencing of 28S rRNA gene

Extraction of host algal DNA was performed on cell pellets from the cultured isolates following a protocol supplied with the DNeasy Plant Kit (Qiagen, Valencia, CA, USA). Polymerase chain reaction (PCR) amplification was performed with 50 μL reaction mixtures containing 23 μL of sterile distilled water, 10 of μL of 5× Phusion HF buffer (New England Biolabs, Beverly, MA, USA), 1 μL of a dNTP mixture (2.5 mM each), 2.5 μL of each primer (10 pmol), 1 μL of Phusion DNA polymerase (2.0 units/50 μL), and 10 μL of template DNA. Partial large subunit (LSU) rDNA sequences were amplified using primers described in Scholin et al. (1994). The ITS-5.8S-ITS2 rDNA sequences were determined using primers described in Nézan et al. (2012). PCR cycling for partial LSU and ITS-5.8S-ITS2 rDNA was carried out in an iCycler Thermal cycler (Bio-Rad, Hercules, CA, USA) using the following conditions: ITS-5.8S-ITS2; pre-denaturation 94°C for 2 min, 37 cycles of 94°C for 30 s, 52°C for 30 s, 72°C for 2 min, and a final extension at 72°C for 5 min, LSU; pre-denaturation 94°C for 3.5 min, 36 cycles of 94°C for 50 s, 45°C for 50 s, 72°C for 80 s, and a final extension at 72°C for 10 min. The PCR products were purified using a QIAquick PCR Purification Kit (Qiagen, Hilden, Germany) according to the manufacturer’s instructions and sequenced (MCLAB, San Francisco, USA). Editing and contig assembly of rDNA sequence fragments were carried out using Bioedit v7.2.5 (Hall, 1999).

### DNA extraction and sequencing of 16S rRNA gene DNA Extraction

For both the lab and field microbiomes, DNA was extracted from filters using a modified CTAB protocol^94^. Briefly, 2 mL microtubes containing membrane filters and an extraction buffer were subjected to three cycles of freezing in liquid nitrogen followed by thawing in a 65°C water bath. Afterward, 8 µL of proteinase K (10 mg mL^-1^ in a TE buffer) was added, and the samples were incubated at 37°C for 30 min. Next, 80 µL of 20% sodium dodecyl sulphate was added, and the samples were incubated at 65°C for 2 h. The solution was then gently mixed with an equal volume of chloroform-isoamyl alcohol (24:1) and centrifuged at 10,000 *g* for 5 min. The aqueous phase was transferred to a fresh 2 mL tube containing 88.8 µL of 3 M sodium acetate (pH 5.1), followed by the addition of 587 µL of isopropanol (≥99%). After centrifugation at 14,000 *g* for 20 min, the supernatant was discarded, and 1 mL of cold 70% ethanol was added. The samples were then centrifuged at 14,000 *g* for 15 min, after which the pellets were air-dried at room temperature and dissolved in 100 µL of TE buffer (10 mM Tris-HCl, 1 mM EDTA, pH 8).

### 16S rRNA Gene Sequencing Lab microbiome

Sequencing libraries were prepared according to Illumina 16S Metagenomic Sequencing Library protocols to amplify the V3 and V4 regions of the 16S rRNA gene. Genomic DNA (2 ng/10 ng input) was amplified by PCR using a 5× reaction buffer, 1 mM dNTP mix, 500 nM each of the universal forward/reverse primers, and Herculase II Fusion DNA Polymerase (Agilent Technologies, Santa Clara, CA, USA). The conditions for the first PCR cycle were 95°C for 3 min for initial heat activation, followed by 25 cycles of 30 s at 95°C, 30 s at 55°C, and 30 s at 72°C, with a final extension at 72°C for 5 min.

The universal primer pair with Illumina adapter overhang sequences used for the initial PCR amplifications were as follows: forward primer (5□-TCGTCGGCAGCGTCAGATGTGTATAAGAGACAGCCTACGGGNGGCWGCAG-3□) and reverse primer (5□-GTCTCGTGGGCTCGGAGATGTGTATAAGAGACAGGACTACHVGGGTATCTAATCC-3□). The first PCR product was purified using AMPure XP beads (Agencourt Bioscience, Beverly, MA, USA). Next, 10 µL of the first PCR product was used for a second round of PCR to construct the final library, incorporating NexteraXT indexed primers. The conditions for the second PCR cycle were identical to the first, except for 10 cycles. The PCR products were again purified with AMPure beads, and the final purified product was quantified using quantitative PCR (KAPA Library Quantification Kit for Illumina Sequencing Platforms) and validated using TapeStation D1000 ScreenTape (Agilent Technologies, Waldbronn, Germany). Sequencing was performed on the Illumina HiSeq platform (Illumina, San Diego, CA, USA).

### Field microbiome

The extracted DNA samples were processed to generate amplicons for variable region 4 (V4) of the nuclear small subunit (SSU) rRNA gene using the universal bacterial/archaeal primers 515F (5′-GTGCCAGCMGCCGCGGTAA-3′) and 806R (5′-GGACTACHVGGGTWTCTAAT-3′), which contain a 6 bp error-correcting barcode unique to each sample. The amplified samples were sequenced on an Illumina MiSeq platform with the 515F and 806R primers.

### Operational taxon unit (microbial lineage) analysis

#### Lab microbiome

The raw data were demultiplexed by index sequences to generate FASTQ files for each sample. Adapter sequences were trimmed using the fastp tool^95^, and error correction was performed for overlapping reads. The paired-end data for each sample were merged into single sequences using FLASH (v1.2.11)^96^. Sequences shorter than 400 bp or longer than 500 bp were discarded. The assembled sequences were processed using QIIME (v1.9)^97^ and UCLUST^98^ for microbial lineage picking in a closed-reference approach. The NCBI 16S Microbial DB was used as the reference. Finally, the microbial lineage abundance table was rarefied to the minimum number of sequences per sample.

#### Field microbiome

The raw sequencing data were processed in R using the EasyAmplicon pipeline package and incorporating various bioinformatics tools^99^. After removing barcodes and primers, paired-end sequences were merged, and quality filtering was applied. Dereplication of full-length sequences was performed, and sequences with a minimum unique size of 10 were retained. MLs were clustered at 97% similarity using de novo chimera removal. Taxonomic classification of the representative microbial lineages was performed against the SILVA 138.2 database^100^ with a confidence threshold of 60%. Finally, the microbial lineage abundance table was rarefied to the minimum number of sequences per sample. The raw sequencing reads were deposited at the NCBI Short Read Archive under BioProject ID PRJNA1180708.

#### Statistical analysis

All statistical analyses were conducted in RStudio (v2024.04.2+764) using R (v.4.3.2). Microbial community comparisons were performed using the phyloseq (v.1.46.0)^101^, Microbiotaprocess (v.1.17.1)^102^, and vegan (v.2.6.6.1)^103^ packages based on ML abundance, taxonomy, and phylogenetic tree data. Visualizations were generated using the ggplot2 package (v.3.5.1) ^104^ as needed. Prior to all analyses, chloroplast-and mitochondria-related microbial lineages were filtered out, and microbial lineage abundance was rarefied. For all analyses except co-occurrence networks, FAPROTAX, and phylosymbiosis, MLs with an average relative abundance of less than 0.005% were excluded. All p-values indicating the significance of microbiome differences between groups were adjusted using the Bonferroni or Benjamini-Hochberg correction. All PERMANOVA permutation tests were conducted with 9,999 permutations. To evaluate the statistical power of detecting group differences in beta diversity via PERMANOVA, a post hoc power analysis was performed using G*Power (v.3.1.9.7)^105^. The analysis was based on a total sample size of 72, with 6 groups, each consisting of 2 strains, and 6 repeated measurements per strain. An effect size (f) of 0.35, an α error probability of 0.05, and a correlation among repeated measures of 0.5 were applied. The power analysis yielded a noncentrality parameter λ of 15.12 and a critical F value of 2.3538, resulting in an achieved power (1-β) of 0.836 (83.6%). These results indicate that the study had adequate statistical power to detect significant between-group differences at a 5% significance level.

Network analysis was carried out using the ggClusterNet R package^106^. Microbial functional prediction was performed using FAPROTAX based on taxonomic data using Python script^107^. To assess the impact of stochastic processes on the microbiome, we applied the normalized stochastic ratio on an NST_jac_ using the NST R package^108^, with host taxonomy (dinoflagellate or diatom) as the grouping variable and strain origin as the meta-group. A correlation analysis of Bray–Curtis dissimilarity between phytoplankton strain microbiomes and the strain’s environmental and spatiotemporal information (Supplementary Table 1) was performed after log-transforming the data and normalizing Euclidean distances between samples to a maximum value of 1. Data for microbiome dissimilarity within the same strain across growth phases and replicates were excluded from the analysis.

### Evolutionary distance and phylosymbiosis analysis

#### Evolutionary distance and host phylogeny

Large subunit ribosomal DNA sequences (total length 1031 bp) were used to calculate evolutionary distances and construct phylogenetic trees among phytoplankton hosts (Supplementary Table 1). For dinoflagellates, LSU sequences from six strains were analysed in this study, with one representative sequence per species chosen to maximize the length of the comparison region (Supplementary Table 4). For diatoms, LSU sequences were downloaded from the NCBI, except for *Coscinodiscus granii*, for which no LSU sequences were available; those of the closely related *Coscinodiscus wailesii* were used instead (Supplementary Table 4). The evolutionary history was inferred using the maximum likelihood method and the Tamura-Nei model^109^. A bootstrap consensus tree based on 500 replicates^110^ was generated to represent the evolutionary history of the analysed taxa. A discrete gamma distribution was used to model evolutionary rate variation among sites (5 categories (+G, parameter = 0.6128)). All evolutionary analyses, including model testing, were performed in MEGA11 ^111^.

#### Microbiota dendrograms

To obtain representative microbiota profiles for each species, the microbial lineage abundance data for each strain and growth phase of the six microalgal hosts were collapsed into a single profile for each species. Beta diversity distances between the six species’ microbiomes were calculated using the Bray–Curtis method. Distance matrices were then clustered using the UPGMA method to generate dendrograms reflecting interspecific microbiome relatedness. To assess the impact of different beta diversity measures on congruence with host phylogeny, dendrograms were generated using weighted UniFrac and unweighted UniFrac distance methods.

#### Robinson–Foulds and matching cluster congruency analyses

Congruence between host phylogeny and microbiota dendrograms was quantified using normalized Robinson– Foulds and normalized matching cluster methods. The analyses were performed using a Python script, with 100,000 random trees used as the comparison baseline^9^.

## Acknowledgements

This research was supported by the National Research Foundation of Korea (NRF) grant funded by the Korea Government (MSIT) (No. RS-2023-00209356 and No. 2022R1C1C1003582), and the Korea Institute of Marine Science & Technology (KIMST) funded by the Ministry of Oceans and Fisheries (RS-2023-00256330, Development of risk managing technology tackling ocean and fisheries crisis around the Korean Peninsula by Kuroshio Current).

## Author contributions

B.S.P. and J.-H.K. conceptualized the project. J.-H.K., J.H.K., Z.L., and Y.K.L. performed the wet laboratory experiments. H.H.S. collected phytoplankton strains. B.S.P., J.-H.K., and J.H.K. collected field phytoplankton bloom samples. J.-H.K. and P.W. conducted the metagenomic analyses. J.-H.K., R.Y.P., and S.H.J. curated the metadata of public samples. J.-H.K., J.H.K., and Z.L. designed the statistical analyses. B.S.P., H.H.S., S.H.B., and M.-S.H. acquired funding. J.-H.K., J.H.K., and Z.L. wrote the manuscript. B.S.P. supervised the work. All authors read and approved the manuscript.

