## Supplementary Figures 1-6 for "Phytoplankton recruitment of specific microbial assemblages and phylosymbiotic patterns"

**File includes Supplementary Figures 1-6.**

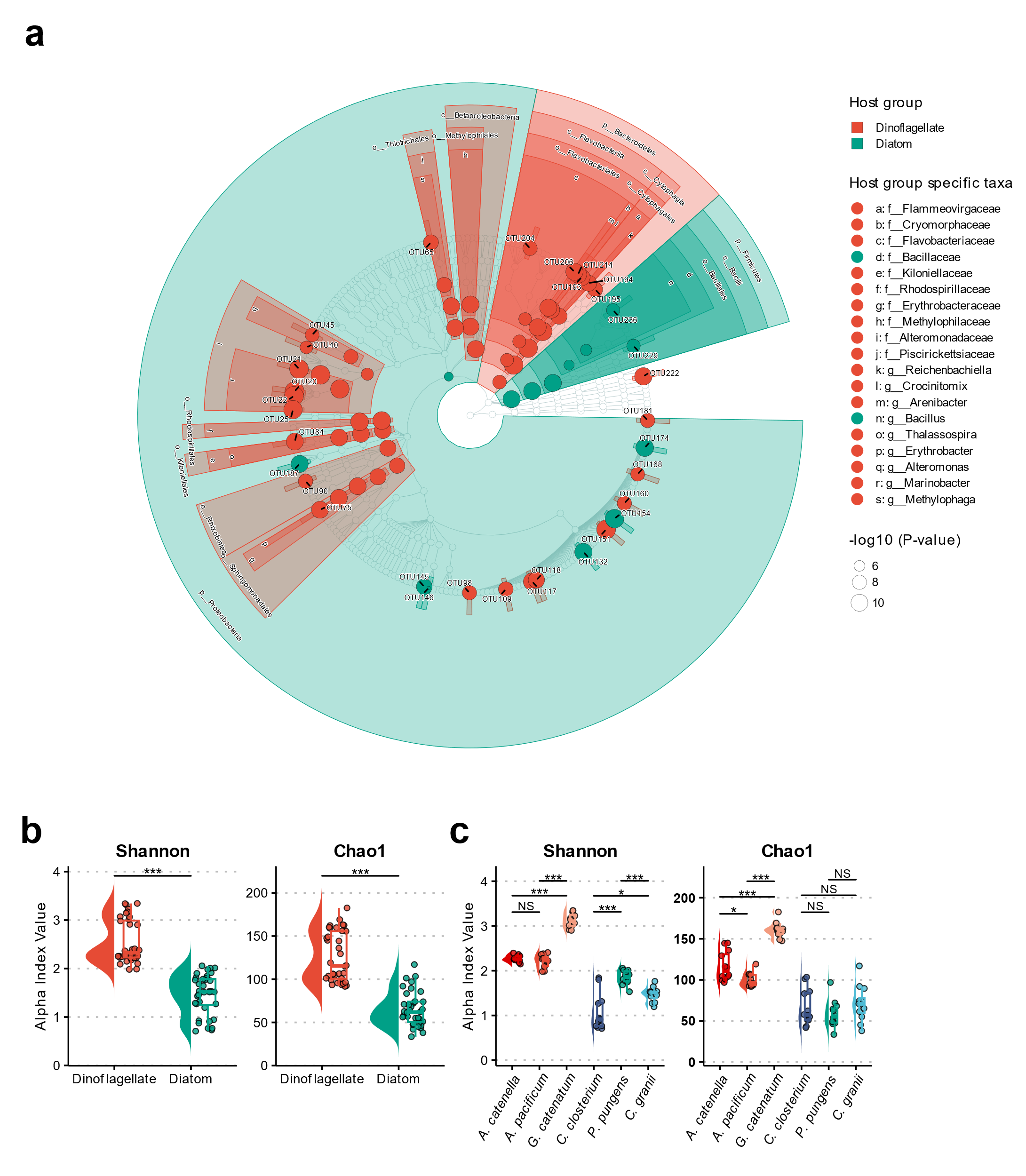

**Supplementary Fig. 1. Significant differences in major bacterial taxa and higher diversity in dinoflagellate microbiomes compared to diatom microbiomes. a**: Microbiome taxa tree illustrating differentially abundant taxa between dinoflagellate and diatom microbiomes. Tree nodes and highlights are colour-coded to indicate the group (dinoflagellate or diatom) in which the corresponding bacterial taxon is significantly more abundant. Node size represents the adjusted p-value (Bonferroni correction). Differential abundance analysis involved a two-step approach: first, a Kruskal–Wallis test was used to identify taxa with overall group differences, followed by a Wilcoxon rank-sum test for pairwise comparisons. Taxa with an absolute linear discriminant analysis) score > 3.12 are shown. **b:** Alpha diversity comparison between dinoflagellate (*n*=36) and diatom (*n*=36) microbiomes. Statistical significance was assessed using a Wilcoxon rank-sum test. **c**: Alpha diversity comparison across six phytoplankton species (*n*=12 per species). Statistical significance was also determined using a Wilcoxon rank-sum test. Significance levels: *P < 0.05, ***P < 0.001; NS: not significant. Boxplot boundaries indicate the first and third quartiles, with the median shown as a line. Whiskers extend to the furthest data points within 1.5× the interquartile range.

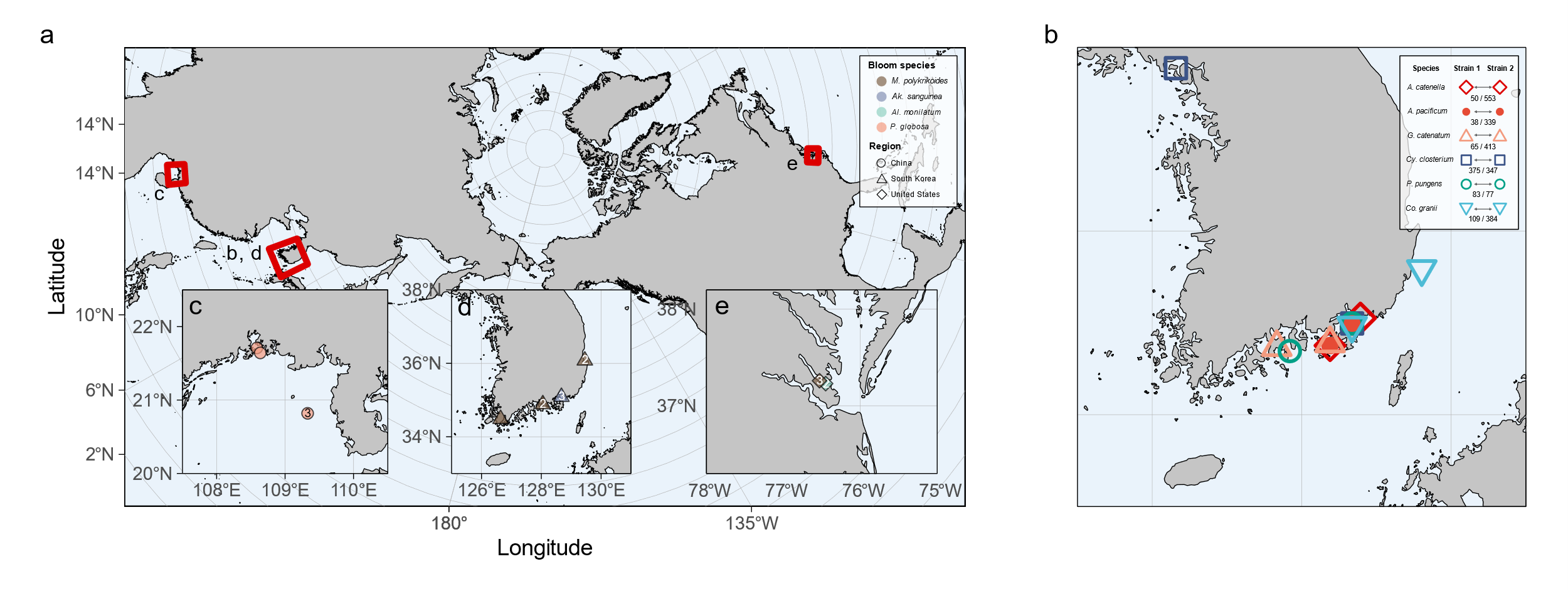

**Supplementary Fig. 2. Geographic locations of host phytoplankton strain isolation and phytoplankton bloom collection analysed in this study. a:** Map showing study regions in Korea, China, and the US. The red rectangle (b, d) represents the area shown in panels b and d, while c and e indicate the regions depicted in panels c and e, respectively. **b:** Isolation sites of phytoplankton strains along the Korean coastline. The numbers shown between the two shapes in the inset (n/n) represent the spatial (km) and temporal (days) distances between the two strains isolated from each location. Supplementary Table 1 provides more details. **c:** Collection sites of phytoplankton blooms in the Beibu Gulf, China. Detailed information can be found in Supplementary Table 3. **d:** Collection sites of phytoplankton blooms along the Korean coastline. **e:** Collection sites of phytoplankton blooms in the York River, eastern US. In panels c to e, the numbers within the symbols indicate the number of samples collected at each site.

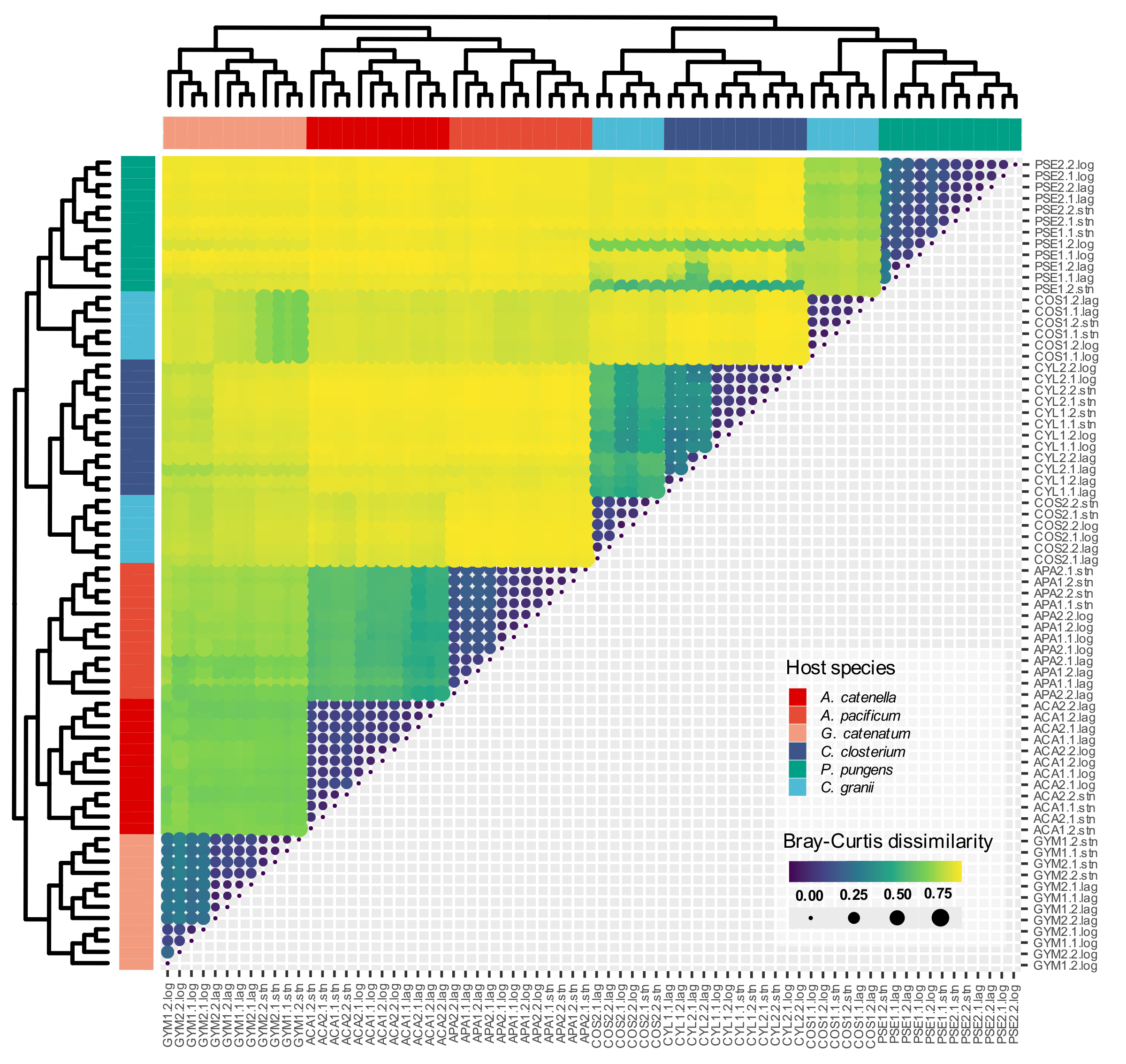
**Supplementary Fig. 3. Heatmap and hierarchical cluster analysis of Bray–Curtis dissimilarity between phytoplankton-associated microbiomes.** The dendrogram was constructed using the UPGMA method. Bray–Curtis dissimilarity is represented by both colour gradation and circle size.

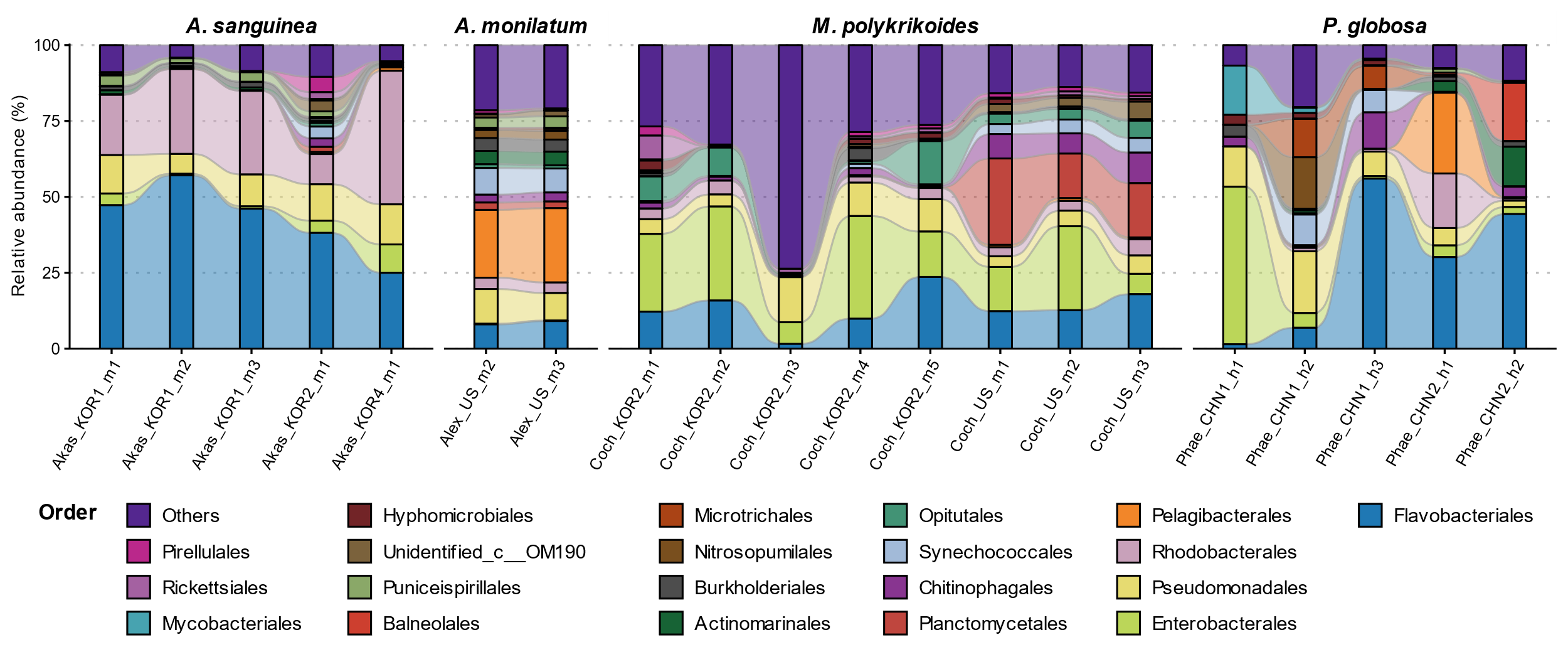

**Supplementary Fig. 4.** **Order-level composition of microbiomes from four microalgal blooms collected across various spatial and temporal scales.** Detailed sample information can be found in Supplementary Table 3.

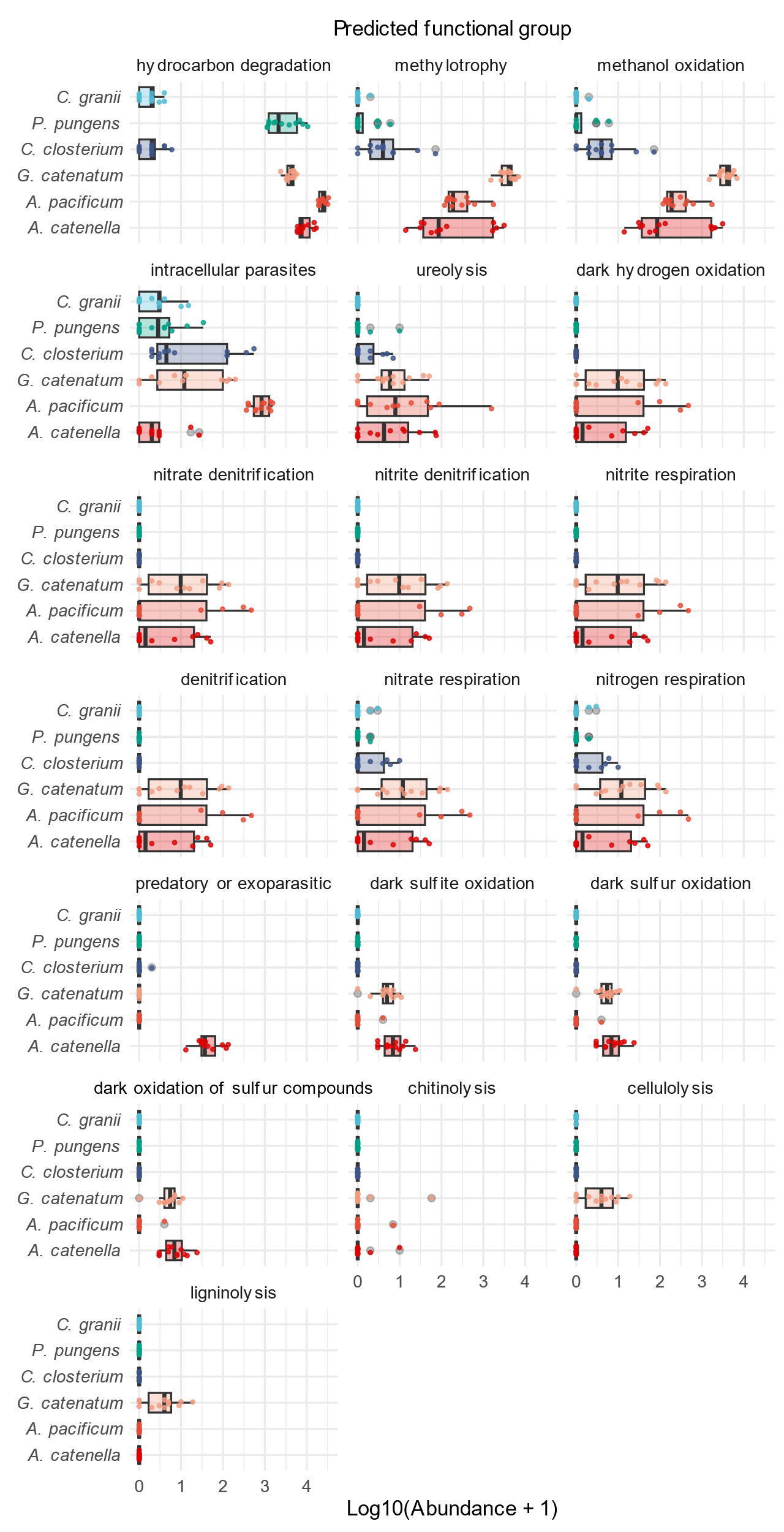

**Supplementary Fig. 5. Distribution of bacterial functional group abundances across different microalgal host species, as predicted by FAPROTAX analysis.** A total of 19 functional groups indicate significant differences between dinoflagellate and diatom hosts, with all groups more abundant in dinoflagellates on average. Panels are arranged in descending order based on linear discriminate analysis scores. Boxplot boundaries indicate the first and third quartiles, with the median shown as a line. Whiskers extend to the furthest data points within 1.5× the interquartile range.

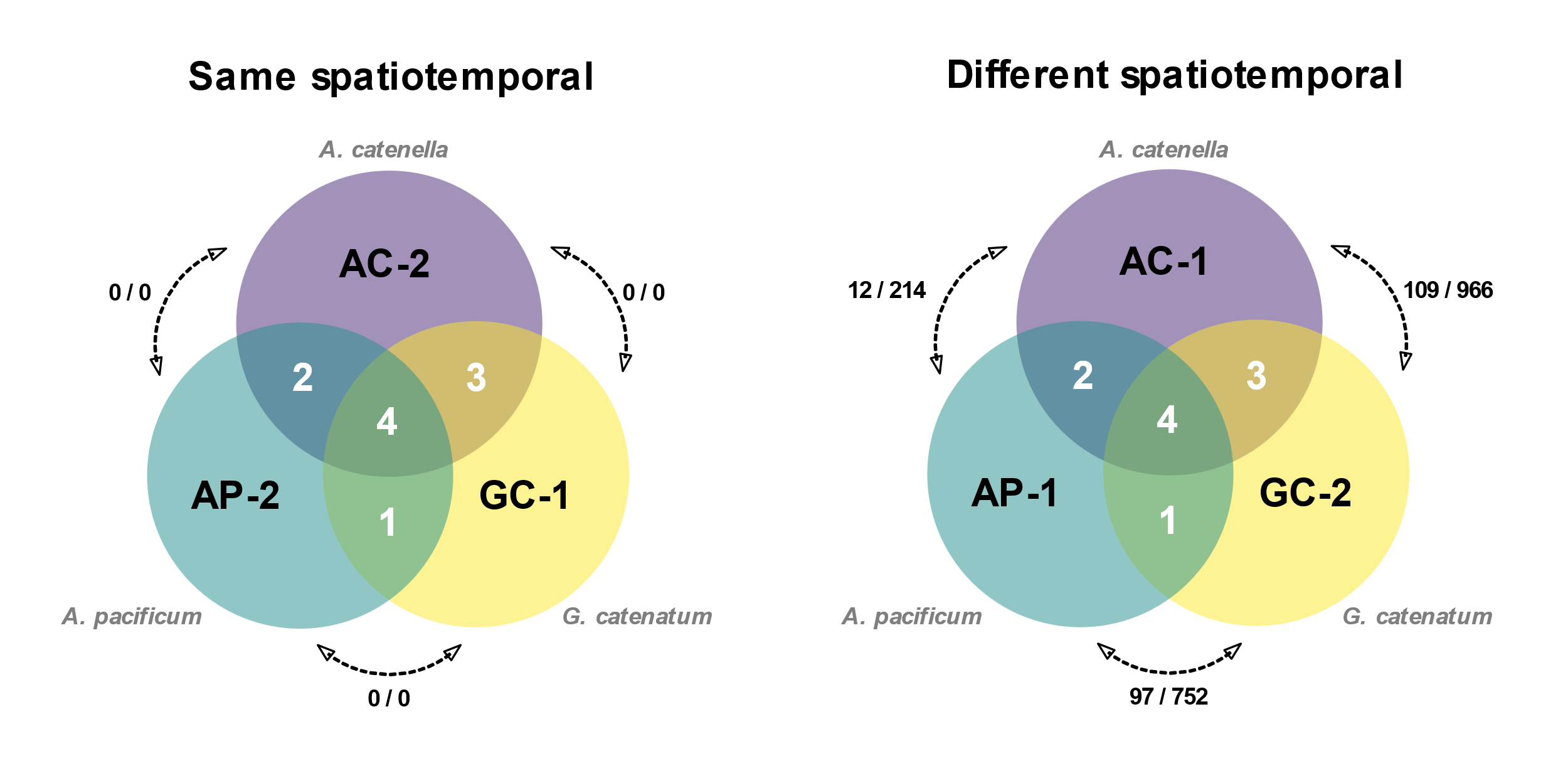
**Supplementary Fig. 6.** **Limited overlap in core bacterial taxa among strains of different species, even when isolated from the same spatiotemporal conditions.** Venn diagrams compare the number of core bacteria (operational taxonomic unit level) shared among three strains of dinoflagellates isolated from the same site on the same day (left) and those isolated from different times and locations (right; details in Supplementary Table 1). Strain names are shown inside the Venn diagram, and species names are labelled outside. The numbers in the intersections represent the shared core bacteria. Arrows between the circles indicate the spatial (km) and temporal (days) distances between the isolation sites of each strain.
